# Metabolomics and proteomics of *L. rhamnosus* GG and *E. coli* Nissle probiotic supernatants identify distinct pathways that mediate growth suppression of antimicrobial-resistant pathogens

**DOI:** 10.1101/2020.12.21.423897

**Authors:** Petronella R. Hove, Nora Jean Nealon, Siu Hung Joshua Chan, Shea M. Boyer, Hannah B. Haberecht, Elizabeth P. Ryan

**Affiliations:** Department of Microbiology, Immunology, and Pathology, College of Veterinary Medicine and Biomedical Sciences; Colorado State University; Department of Environmental and Radiological Health Sciences, Colorado State University. Fort Collins, Colorado. 80523.; Department of Chemical and Biological Engineering. Colorado State University, Fort Collins, Colorado. 80523

**Author notes:** Correspondence: Elizabeth P. Ryan, PhD Associate Professor Department of Environmental and Radiological Health Sciences College of Veterinary Medicine and Biomedical Sciences Colorado State University/Colorado School of Public Health Fort Collins, CO 80524. Petronella R. Hove* and *Nora Jean Nealon* contributed equally to this work.

**Keywords:** Antimicrobial resistance, probiotics, cell free, supernatants, growth suppression, proteome, metabolome

## Abstract

Probiotics merit testing as alternatives to conventional antibiotics and are receiving increased attention for efficacy against multi-drug resistant pathogen infections. This study hypothesis was that the Gram-positive probiotic, *L. rhamnosus* GG (LGG) and Gram-negative *E. coli* Nissle (ECN) secrete distinct proteins and metabolites to suppress pathogen growth. LGG and ECN cell free supernatants were tested in a dose-dependent manner for differential growth suppression of *Salmonella* Typhimurium, *Escherichia coli*, and *Klebsiella oxytoca* that harbor antimicrobial resistance (AMR). Across supernatant doses, LGG was 6.27% to 20.55% more effective than ECN at suppressing AMR pathogen growth. Proteomics and metabolomics were performed to identify pathways that distinguished LGG and ECN for antimicrobial functions. From the 667 detected metabolites in probiotic cell free supernatants, 304 metabolites had significantly different relative abundance between LGG and ECN, and only 5 and 6 unique metabolites were identified for LGG and ECN respectively. LGG and ECN differences involved amino acid, energy and nucleotide metabolism. Proteomics analysis of ECN and LGG cell free supernatants identified distinctions in 87 proteins, where many were related to carbohydrate and energy metabolism. Integration of genome-proteome-metabolome signatures from LGG and ECN with predictive metabolic modeling supported differential use of substrates by these two probiotics as drivers of antimicrobial actions. ECN metabolized a range of carbon sources, largely purines, whereas LGG consumed primarily carbohydrates. Understanding functional biosynthesis, utilization and secretion of bioactive metabolites and proteins from genetically distinct probiotics will guide strategic approaches for developing antibiotic alternatives and for controlling spread of multi-drug resistant pathogens.

**Importance:** Probiotics are practical alternatives for protection against antimicrobial resistant pathogens. Bioactive probiotics molecules merit further investigation using high throughput - omic approaches. This study identified functional differences between Gram-positive *L. rhamnosus* GG (LGG) and Gram-negative *E. coli* Nissle (ECN) probiotics that suppressed the growth of antimicrobial resistant *S.* Typhimurium, *K. oxytoca*, and *E. coli*. Proteomes and metabolomes of the probiotic cell free supernatants showed metabolic differences between LGG and ECN for mediating pathogen growth suppression. Metabolites distinguishing LGG versus ECN growth suppression included carbohydrates, lipids, amino acids, and nucleic acids. The metabolic flux differences between ECN and LGG, which coincided with observed separations in the proteomes and metabolomes, was hypothesized to explain the differential suppression of AMR pathogens. Integrated metabolite and protein signatures produced by each probiotic merit attention as adjuvant therapeutics for antimicrobial resistant infections.

## Introduction

Probiotic microorganisms have been largely explored for their capacities to suppress pathogen growth. Conventional paradigms consider the broad-acting mechanisms by which probiotics antagonize pathogens, such as competitive exclusion of pathogens in host tissues, production of organic acids, and modulation of host immunity [1]. Few studies exist for comparison of probiotics, although there is evidence that they have species and strain-dependent differences in **AMR** pathogen growth suppression [1–6]. According to the World Health Organization and Centers for Disease Control, the burden of AMR infections from Enterobacteriaceae accounts for ~423,000 infections annually and adds ~$1.5 billion to healthcare costs [7, 8]. Among AMR Enterobacteriaceae, the non-typhoidal *Salmonella* and most frequently *S*. Typhimurium, is a key global cause of diarrheal diseases that accounts for approximately 1.2 million illnesses each year in the United States alone [9]. A large body of research has characterized the spread of AMR *E. coli* isolated from people, livestock, and in environmental waters, and there are reports of human clinical infections with *E. coli* susceptible to last-resort antimicrobial agents, including colistin [10, 11]. *Klebsiella* species, including *K. pneumoniae* and *K. oxytoca* are also part of the Enterobacteriaceae family. These species are normally present in environmental samples and notable for causing hospital and community-acquired opportunistic infections in the urinary tract, respiratory tract, and bloodstream [12]. *Klebsiella* species readily form biofilms [13] that contributes to innate AMR, and making them particularly difficult to eradicate [12, 14].

Gram-positive probiotic *Lactobacillus rhamnosus* GG (**LGG**) and the Gram-negative probiotic *Escherichia coli* Nissle (**ECN**) have been well-characterized in their capacities to antagonize antimicrobial-resistant Enterobacteriaceae [15, 16]. Recent studies illustrate that they differentially regulate host gut immunity to protect against enteric infections [17], suggesting that they may also function to antagonize pathogens through distinct mechanisms. Additional investigations have identified bioactive small molecules, including multiple amino acids, carbohydrates and lipids, that are produced by LGG and ECN that contribute to their anti-pathogen activities [18–21]. Other investigations have identified proteome and genome markers in LGG and ECN including genes that regulate the production of lipids, amino acids, and energy metabolites for anti-cancer, anti-inflammatory, and pathogen-protective capacities [19, 21, 22], suggesting that these chemical classes are important small molecule contributors to their antimicrobial activity.

This study was designed to examine and establish mechanisms by which the cell free supernatants from Gram-positive and Gram-negative probiotics, namely LGG and ECN, differentially function to suppress AMR pathogens, *E. coli*, *S.* Typhimurium, and *K. oxytoca*. This study hypothesis was that differential nutrient metabolism by LGG and ECN leads to production of distinct antimicrobials that exhibit dose-dependent differences in growth suppression of antimicrobial resistant pathogens, namely *S*. Typhimurium, *Klebsiella*, and *E. coli*. The composition of cell free supernatants from Gram-positive and Gram-negative probiotics, namely LGG and ECN, that suppress *E. coli*, *S.* Typhimurium, and *K. oxytoca* growth was assessed by an integrated, non-targeted metabolomics, proteomics, and metabolic network analysis. Metabolic differences between a Gram-positive and Gram-negative probiotic for antimicrobial functions represents a novel approach with broad-spectrum applications to environmental, animal and human health.

## Results

### Phenotypic and genotypic characterization of three AMR pathogens

*E. coli*, *S.* Typhimurium, and *K. oxytoca* pathogens were screened for phenotypic AMR against five representative drug classes using Kirby-Bauer disk diffusion (**Table 1**). All three pathogens were resistant to the beta-lactam drugs ampicillin and cefazolin. In addition, *S.* Typhimurium displayed multidrug resistance by displaying additional resistance to the aminoglycoside drug gentamicin as well as tetracycline. To further characterize AMR genes by *E. coli*, *S.* Typhimurium, and *K. oxytoca,* the genomes were **BLAST**(Basic Local Alignment Search Tool)-searched against a routinely curated AMR gene database (**Fig. 1**). One hundred and twelve antimicrobial genes spanning 15 functional classes were identified across the three pathogens (**Fig. 1** & **File S1**). Many of the genes encoded multidrug resistance efflux pumps and efflux pump regulators across the three pathogens and were followed by beta-lactam resistance genes including class A, B and C beta lactamases and penicillin binding proteins. Gene classes that distinguished the three pathogens were multi-drug resistance ribosomal target modifiers detected exclusively in *S.* Typhimurium, tetracyclines detected only in *E. coli*, and phenicols, rifampins and regulator proteins identified exclusively in the *K. oxytoca* genome.

**Table 1.** Kirby-Bauer disk diffusion for antimicrobial resistant pathogens

|  | *S. Typhimurium* | *E. coli* | *K. oxytoca* |
| --- | --- | --- | --- |
| Antimicrobial Agent | Zone Diameter (mm), (Designation) |  |  |
| Amikacin (AK-30) | 26.9 ±2.3, (S) | 24.3±2.3, (S) | 31.3±2.1, (S) |
| Ampicillin (AMP-10) | 6.0 ± 0.0, (R) | 6.0 ± 0.0, (R) | 0.0±0.0, (R) |
| Cefazolin (CZ-30) | 6.0 ± 0.0, (R) | 7.8±4.5, (R) | 14.2±9.6, (R) |
| Ciprofloxacin (CIP-5) | 26.3±3.8, (S) | 25.8±5.3, (S) | 22.9±4.4, (I) |
| Gentamicin (CN-10) | 7.1±3.2, (R) | 20.2±2.7, (S) | 18.5±3.3, (S) |
| Linezolid (LZD-30) | 6.0 ± 0.0, (NA) | 6.2±0.4, (NA) | 1.8±2.6, (NA) |
| Meropenem (MEM-10) | 32.6 ± 0.79, (S) | 34.3±1.0, (S) | 40.2±2.5, (S) |
| Penicillin (P-10) | 6.0 ± 0.0, (NA) | 8.5±2.5, (NA) | 0.8±3.4, (NA) |
| Tobramycin (NN-10) | 17.4 ±3.9, (S) | 23.7±1.2, (S) | 31.0±2.4, (S) |
| Tetracycline (TE-30) | 9.6 ±7.0, (R) | 17.2±4.0, (I) | 28.5±3.1, (S) |
| Vancomycin (VA-30) | 6.3 ±0.76, (NA) | 6.7±0.8, (NA) | 0±0.0, (NA) |

Values represented mean ± standard deviation of the zone of inhibition diameter measured in millimeters. Designations are defined as Susceptible “S”, Intermediate “I” or Resistant “R” based on standards defined by the Clinical Laboratory and Standards Institute (CLSI) for
*Enterobacteriaceae* species.
“NA” indicates cutoff value for “S”, “I”, or “R” not defined by the CLSI.

**Figure 1.**
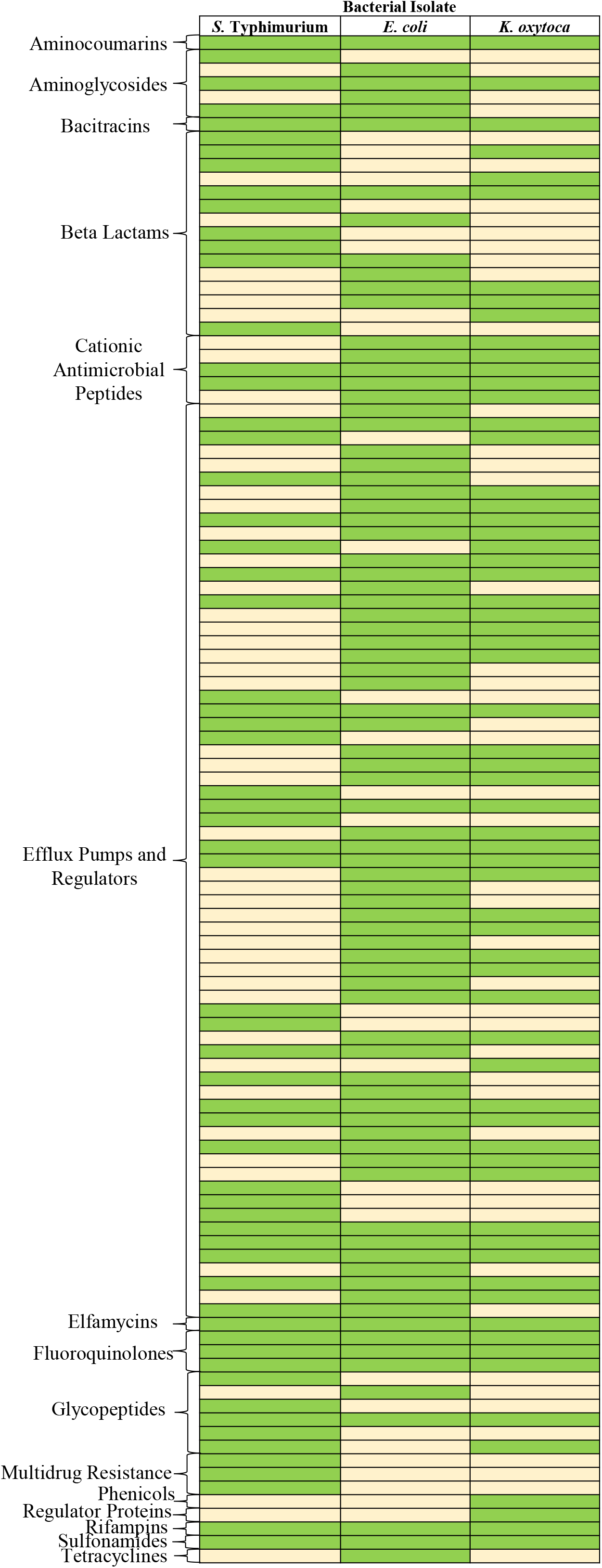
AMR genes identified in the pathogen isolate genomes *E. coli*, *S. Typhimurium*, and *K. oxytoca*. Green boxes indicate gene presence while tan boxes indicate gene absence. Approximately 112 antimicrobial resistance genes spanning 15 functional classes were identified across the three pathogens.

### Differential AMR pathogen growth suppression by L. rhamnosus GG and E. coli Nissle cell free supernatants

LGG and ECN supernatants dose dependently (12% v/v to 25% v/v) decreased AMR *S.* Typhimurium, *E. coli*, and *K. oxytoca growth*. **Fig. 2** shows the minimum inhibitory cell free supernatant dose-response for each pathogen, defined as the dose of supernatant that enhanced pathogen growth suppression compared to the vehicle control. Next, the percent growth inhibition was calculated for each probiotic supernatant relative to the vehicle control. Across all three AMR pathogens and for each probiotic supernatant concentration, LGG was 6.27% to 20.55% more effective at suppressing pathogen growth when compared to ECN (**Fig. 2**). **Fig. S1** shows the quantification and comparisons for ECN and LGG growth suppression for each pathogen at 4h intervals.

**Figure 2.**
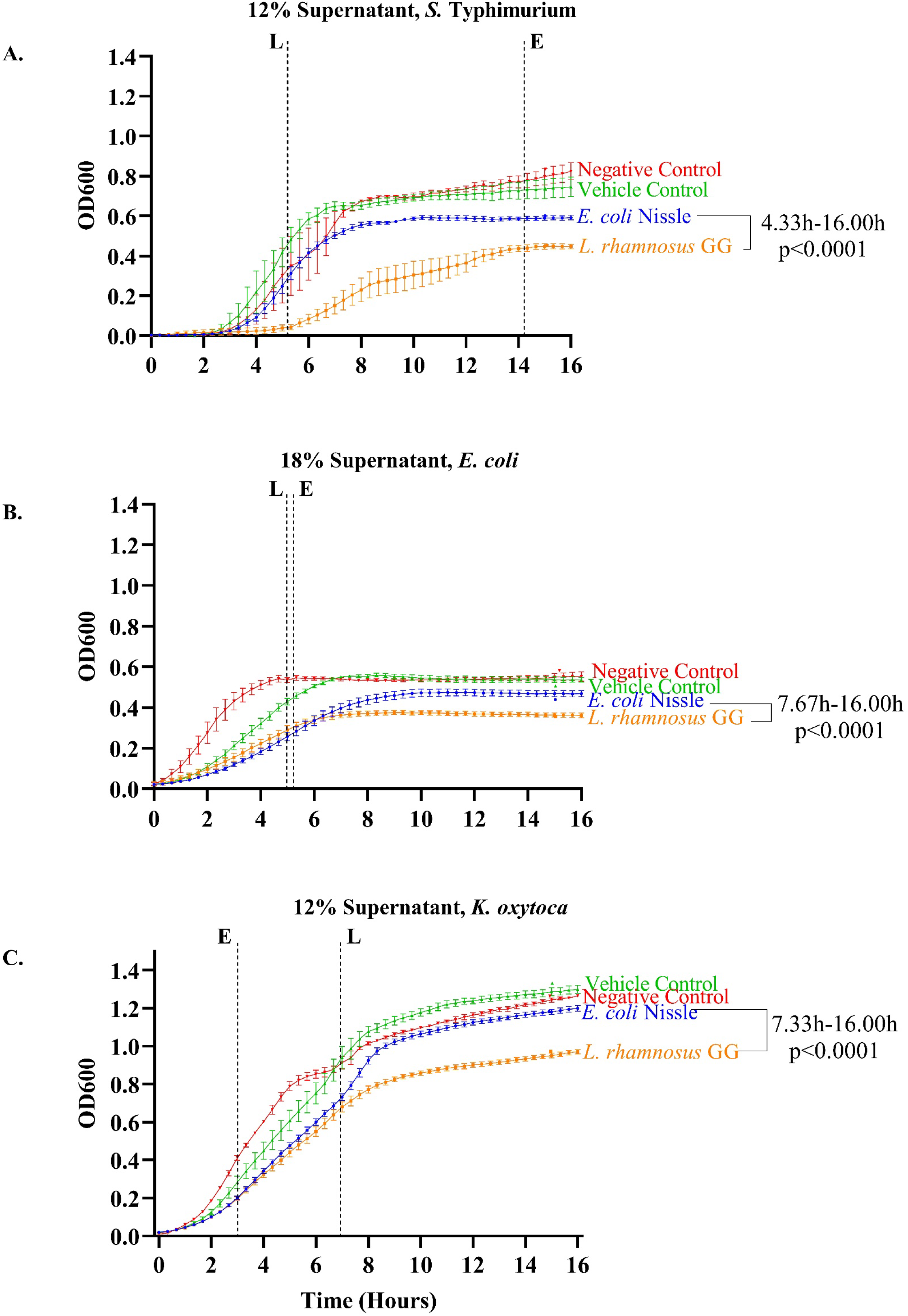
AMR pathogen growth suppression by ECN and LGG probiotic cell free supernatants. Figures depict the growth curves of *S.* Typhimurium, *E. coli* and *K. oxytoca* recorded over 18 hours under the minimum inhibitory dose (supernatant volume/total volume *100) of probiotic cell free supernatant. Bacterial abundance is reported through optical density readings at a wavelength of 600 nm (OD600). The minimum supernatant doses at which both *L. rhamnosus* GG (LGG) and *E. coli* Nissle (ECN) supernatants achieved growth suppression for *S. Typhimurium*, *E. coli* and *K. oxytoca* were the 12%, 18% and 12% respectively. Maximal *Salmonella* growth suppression was achieved at 5.33h for *L. rhamnosus* GG (41.20% p<0.0001) (Dashed line-L) and at 13.67h for *E. coli* Nissle (11.48%, p <0.01) (Dashed line-E). For pathogenic *E. coli* maximum growth suppression for LGG supernatant was 30.40% and occurred at 4.67h (p<0.0001). For the *E. coli Nissle* supernatant, maximal pathogenic growth suppression occurred at 5.00h at 29.45% (p<0.0001). *L. rhamnosus* GG suppressed *K. oxytoc*a growth between 3.00h-16.00h and achieved a maximal percent growth of 28.85% suppression at 7.33h (p<0.0001). *E. coli* Nissle suppressed *K. oxytoca* growth between 3.00h-16.00h and reached maximal growth suppression of 23.86% at 3.33h (p = 0.0035). Dashed lines indicate maximum growth suppression observed for LGG (black) or ECN (blue).

The 12% v/v was the minimum supernatant dose at which both LGG and ECN supernatants achieved growth suppression of *S.* Typhimurium (**Fig. 2A)**. LGG suppressed *S.* Typhimurium growth between 4.33h-16.00h (p<0.0001), and ECN between 13.33h-16.00h (p<0.05) when compared to the vehicle control treatment. Maximal *S.* Typhimurium growth suppression was achieved at 5.33h for LGG (41.20%, p<0.0001) and at 13.67h for ECN (11.48%, p<0.01). LGG supernatant was 6.27% more effective at suppressing *S.* Typhimurium growth compared to ECN (p<0.0001, 8.33h).

The 18% v/v supernatant dose was the lowest dose where LGG and ECN supernatant suppressed *E. coli* growth compared to the vehicle control (**Fig. 2B)**. At this dose, LGG supernatant suppressed *E. coli* growth 30.40% more than the vehicle control (p<0.0001, 4.67h) and ECN supernatant was 29.45% more effective than the vehicle control (p<0.0001, 5.00h). When comparing probiotic supernatants, LGG was 20.55% more effective than ECN at suppressing *E. coli* growth (p<0.0001, 16.00h).

For *K. oxytoca*, the 12% v/v supernatant was the lowest dose where LGG and ECN achieved growth suppression versus the vehicle control (**Fig. 2C)**. At this dose, LGG suppressed *K. oxytoc*a growth between 3.00h-16.00h and achieved a maximal percent growth of 28.85% suppression at 7.33h (p<0.0001). ECN suppressed *K. oxytoca* growth earlier than LGG, between 3.00h-16.00h and reached maximal growth suppression of 23.86% at 3.33h (p<0.005). At maximal growth suppression, LGG was 19.30% more effective than ECN at suppressing *K. oxytoca* growth (p<0.0001, 11.00h).

### E. coli Nissle and L. rhamnosus GG cell free supernatants exhibit differences in metabolic pathways and metabolite production

Given that LGG supernatant was more effective at suppressing growth of all three AMR pathogens tested when compared to ECN supernatant (**Fig. 2**), we next applied a global, non-targeted metabolomics analysis to the probiotic cell free supernatants. A total of 667 metabolites were detected in the LGG and ECN supernatant metabolomes, and the major metabolomic differences between LGG and ECN are depicted in **Fig. 3**. The complete metabolome is provided in **File S2**. Among the 667 detected metabolites, 412 metabolites were characterized: 155 amino acids, 28 carbohydrates, 10 energy metabolites, 63 lipids, 61 nucleotides, 44 xenobiotics/other metabolites, 32 peptides, and 19 cofactors and vitamins. Two-hundred and fifty-five metabolites were unnamed and reported by their mass to charge ratio (m/z) and retention index (RI). Of the 667 total metabolites detected, 304 were differentially abundant (p<0.05) between LGG and ECN with only 5 and 6 unique metabolites identified respectively. **Fig. 3A** shows common, shared, and unique metabolites among ECN, LGG, and MRS broth. Principal coordinates analysis for metabolite median-scaled abundances showed a clear separation of ECN versus LGG metabolomes (**Fig. 3B**). A complete list of metabolites with statistically different median-scaled abundances is provided in **Table S1**. **Fig. 3C** shows the top 50 metabolites ranked according to statistical p-value differences between LGG and ECN, where the median-scaled abundance in fold-difference between ECN versus LGG is depicted. A complete list of differentially abundant metabolites between LGG versus ECN supernatants are reported in **Table S1**.

**Figure 3.**
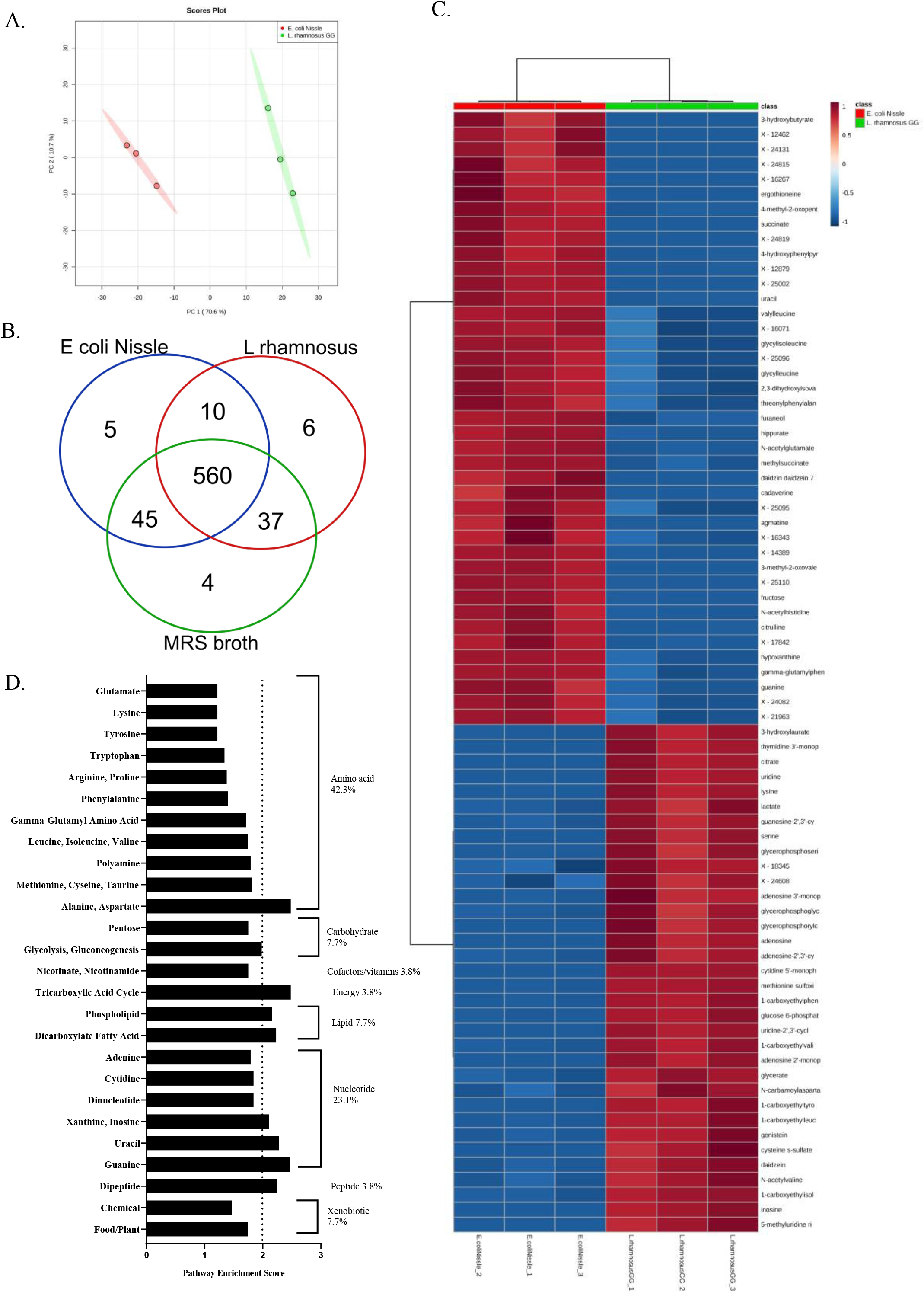
Global, non-targeted metabolomes of *L. rhamnosus* GG and *E. coli* Nissle cell-free supernatant. **A**. Principal component analysis of *L. rhamnosus* GG (LGG) and *E. coli* Nissle (ECN) supernatant and vehicle control media. Each circle represents a biological replicate. **B**. Venn diagram illustrating metabolite presence versus absence differences in ECN versus LGG along with metabolites not present in the vehicle control (MRS broth) when compared to probiotic supernatants. **C**. Heat map of 50 metabolites ranked according to magnitude of fold-differences between ECN and LGG. **D.** Pathway enrichment scores for metabolic pathways that contributed to significantly different metabolites when comparing ECN versus LGG.

Pathway enrichment scores (**PES**) were calculated to evaluate the contribution of metabolic pathways to metabolites with significantly different abundances for ECN versus LGG **Fig. 3D**. Amino acids accounted for 42.3% of metabolite profile differences between ECN and LGG. Polyamine metabolism (PES 1.10) contained the metabolite cadaverine (37.46 -fold higher in ECN versus LGG supernatant, p<1.00E-30), which has been reported to enhance the effectiveness of carboxypenicillins against *P. aeruginosa* [23]. Agmatine (11.35-fold higher in ECN versus LGG supernatant) also has some antiviral activity [24] and antimalarial effects [25] as well as antibacterial effects [26, 27]. Methionine, cysteine, S-adenosyl methionine and taurine metabolism also distinguished ECN versus LGG metabolism with methionine sulfone (0.36-fold lower in ECN vs LGG, p<0.001) and methionine sulfoxide (0.090-fold lower in ECN vs LGG, p<1.00E-30). Methionine sulfoxide has been reported to enhance penicillin susceptibility of highly refractory Gram negative organisms [28] and methionine sulfone was shown to impair glutamate and methionine metabolism in *Salmonella*, *Klebsiella*, and other pathogens [29, 30].

Nucleotides accounted for 23.1% of metabolic pathway differences when comparing ECN to LGG with xanthine, inosine (PES 2.11), uracil (PES 2.80) and guanine metabolism (PES 2.47). Hypoxanthine was 5.17-fold higher in ECN versus LGG supernatant, and whose oxidation to xanthine has been shown to produce antibacterial reactive species [31]. Carbohydrate metabolism contributing to metabolite profile differences between ECN and LGG involved glycolysis and gluconeogenesis (PES 1.21). Glycolytic metabolites included glucose 6-phosphate (0.060 fold lower in ECN versus LGG, p<1.00E-30), and lactate (0.31 fold lower in ECN versus LGG, p<1.00E-10), which have shown direct bactericidal effects on Gram-negative bacteria [32].

In addition to metabolites that were differentially abundant between ECN and LGG, metabolome analysis also revealed distinct metabolites from MRS broth that are depleted or accumulated only in LGG or ECN supernatant respectively, (**File S3**).

### Distinct proteome compositions for E. coli Nissle and L. rhamnosus GG supernatants

The non-targeted proteome of ECN and LGG cell-free supernatants was explored for mechanistic contributions to AMR pathogen growth suppression (**Fig. 4**). The complete probiotic cell free supernatant proteome, protein accession numbers, and gene-ontology terms are provided in **File S4** for 130 total proteins. Forty-nine of these proteins were from animal origin as arising from culture media-broth and were excluded from downstream analysis. Of the remaining proteins, 67 had ECN origin and 14 proteins had LGG origin specificity and only one protein, glyceraldehyde 3-phosphate dehydrogenase, was identified in both ECN and LGG supernatants (**Fig. 4A**).

**Figure 4.**
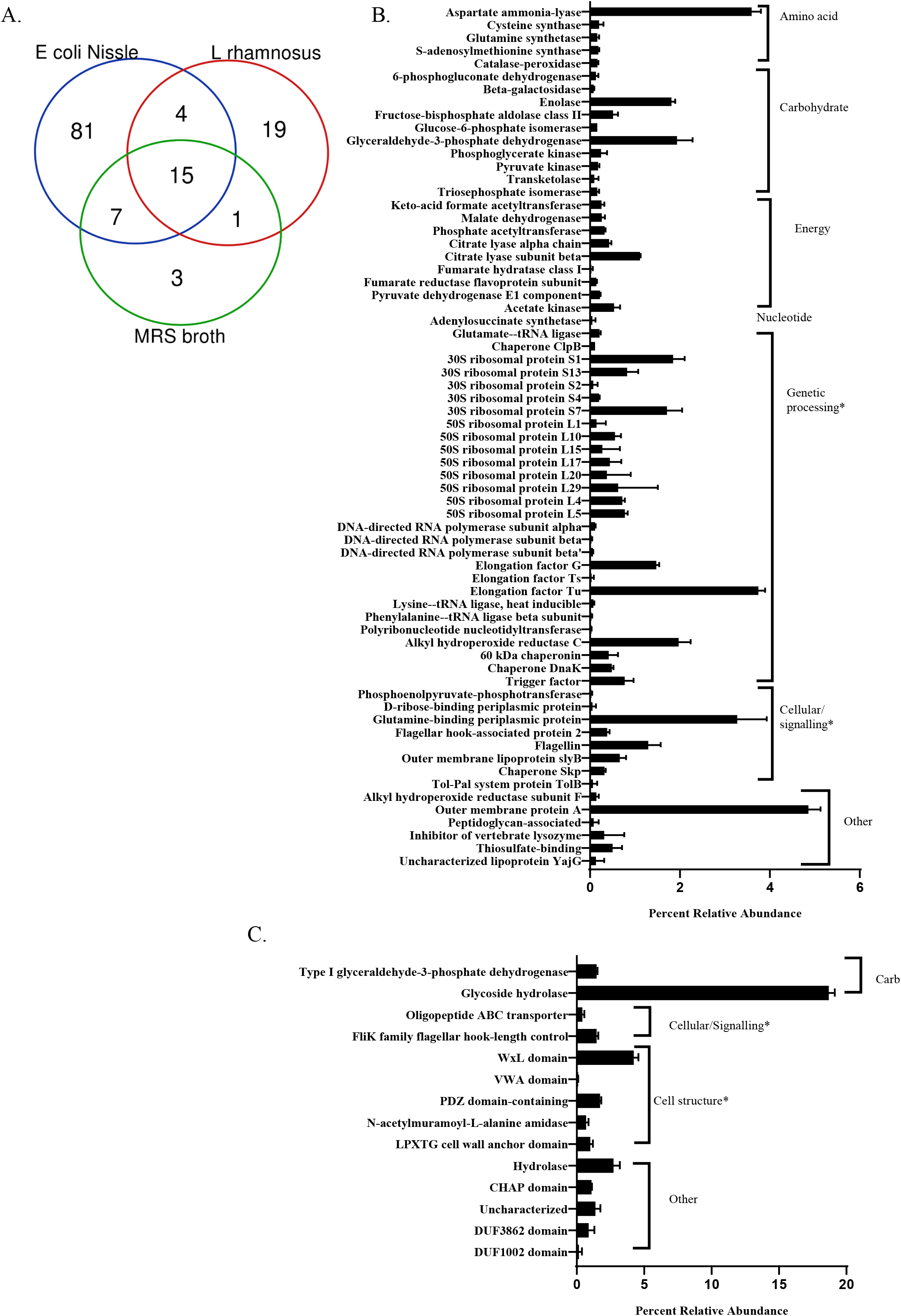
Non-targeted proteome of *E. coli* Nissle and *L. rhamnosus* GG cell free supernatants. **A**. Venn diagram shows the number of proteins identified in *E. coli* Nissle (ECN) supernatant, *L. rhamnosus* LGG supernatant, and sterile MRS broth. Percent relative abundances of proteins found in cell free supernatants of **B**. *L. rhamnosus* GG supernatant and **C**. *E. coli* Nissle supernatant.

The LGG supernatant proteome is classified in **Fig. 4B**. Notably, the glycolysis enzyme glyceraldehyde 3-phopshate dehydrogenase represented 1.47% of the LGG supernatant proteome, Other proteins identified included the CHAP (cysteine and histidine-dependent aminohydrase/proteases) and at hydrolase domain protein (2.73%). The remaining 3 proteins detected in LGG supernatant were uncharacterized. In the ECN supernatant-proteome, proteins involved in carbohydrate metabolism (14 proteins) and amino acid metabolism (8 proteins) were identified (**Fig. 4C**). Among carbohydrate metabolism proteins, the glycolysis enzyme glyceraldehyde 3-phosphate (1.93% abundance) was the most elevated, whereas contributions from enolase (1.81%), another glycolytic protein, were also observed. Major contributors to amino acid metabolism included the aspartate metabolism enzyme aspartate ammonia lyase (3.59 % of total proteome abundance) and glutamine-binding periplasmic protein (3.28% abundance) that is responsible for glutamine transport.

### Metabolic modeling predicts metabolites and proteins contributing to pathogen growth suppression using E. coli Nissle and L. rhamnosus supernatants

Under the assumption that depletion and accumulation of metabolites identified in the cell free supernatants were caused by metabolite consumption and production by the probiotics, metabolic modeling was used to explore the differential dependencies of ECN and LGG on metabolites and associated metabolic enzymes for ATP production (i.e. growth promotion). **Fig. 5** shows the simulated flux distributions for LGG and ECN, which reflect their differential use of carbon sources. Flux distributions were simulated by performing parsimonious flux balance analysis (**pFBA**) on draft metabolic models for the two probiotics (reconstructed using KBase) constrained by the relative metabolite consumption and production observed in the metabolomics data. Of the 667 metabolites detected in the supernatant metabolome, 204 metabolites were present in at least one of the reconstructed models.

**Figure 5.**
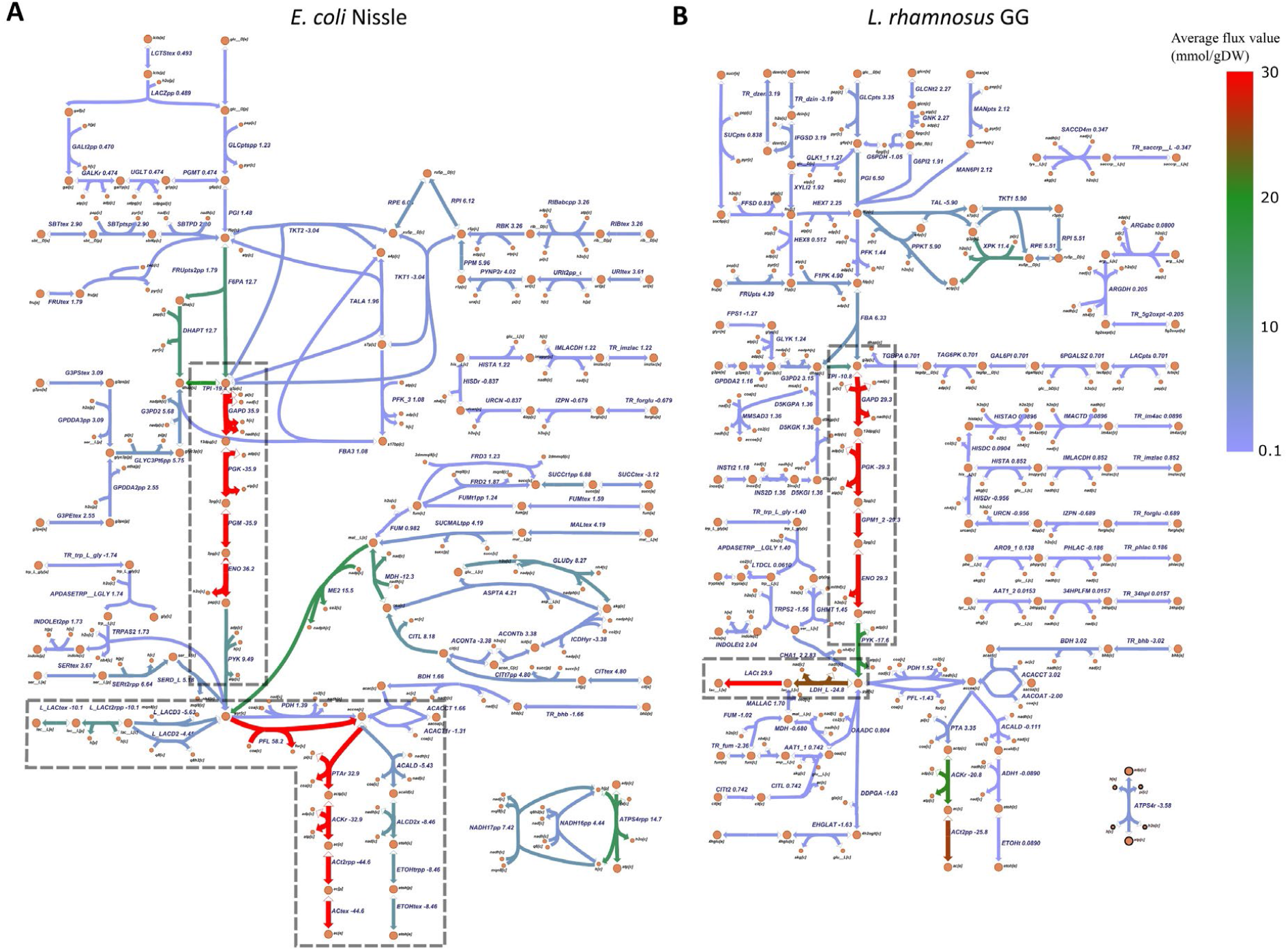
Predicted metabolism of **(A)** *E. coli* Nissle (ECN) and **(B)** *L. rhamnosus* GG (LGG) by parsimonious Flux Balance Analysis (**pFBA**) under the constraints of relative consumption and production of metabolites inferred from the metabolomics dataset. The flux values shown are the average values of 10,000 simulations, normalized by the biomass production, in the unit of mmol / gram cell dry weight. The color of each reaction changes with the magnitude of the average flux as shown in the color bar. The entire dataset is available as **File S5**.

The ECN supernatant-metabolite profiles revealed the capacity for ECN to metabolize a range of carbon sources, such as sugar, nucleoside, amino acid, glycerophospholipid, whereas LGG consumes primarily carbohydrates. Both organisms rely on the lower part of glycolysis for ATP generation, which was supported by the presence of these enzymes in the proteome data (**Fig. 4**). Under the microaerobic conditions under which the probiotic cultures were maintained, LGG relies primarily on lactate production, and was confirmed by the significantly increased lactate detected in LGG supernatant versus ECN (0.31, p<1.00E-13). The heterofermentive probiotic ECN-pFBA predicted that ECN can use lactate, ethanol, and succinate that are produced as terminal electron acceptor for anaerobic respiration and ATP generation by ATP synthase which produced fluxes in 99.79% of the ECN flux samples. Consistent with this flux analysis finding is that the ECN supernatant metabolome had a significantly higher succinate abundance (8.98, p<1.00E-30).

Given the metabolite consumption, biosynthesis profiles and the simulated flux distributions, shadow price (**File S5**) was performed to represent the increase/decrease in the objective function value (i.e. probiotics growth) per unit of increase in resource available reflected in a constraint (i.e. the required consumption/production of metabolites). Based on this rationale, the growth-promoting/competing role of each metabolite that was detected in the metabolome and is present in the models was analyzed.

## Discussion

The differential efficacy of cell free supernatants from two distinct probiotics were investigated for AMR pathogen growth suppression. The gram-positive probiotic LGG suppressed growth of three AMR pathogens, *S.* Typhimurium, *E. coli*, and *K. oxytoca* with lower doses and exposure time when compared to the Gram-negative probiotic ECN. These pathogens collectively contained and expressed resistance to multiple antimicrobial drug classes, emphasizing the need to identify targeted solutions for suppressing growth. To evaluate and compare the small molecule contributors to the differential antimicrobial activity of ECN versus LGG cell-free supernatant, a global, non-targeted metabolomics and proteomics analysis was applied. The proteomes and metabolomes of each probiotic supernatant was integrated with the genomes to develop predictive metabolic models. This integrated multi-omic systems modeling approach predicted major metabolic differences influencing the composition of ECN and LGG supernatants, namely differential regulation of carbohydrate, energy, nucleotide, and amino acid pathways (**Fig. 6**).

**Figure 6.**
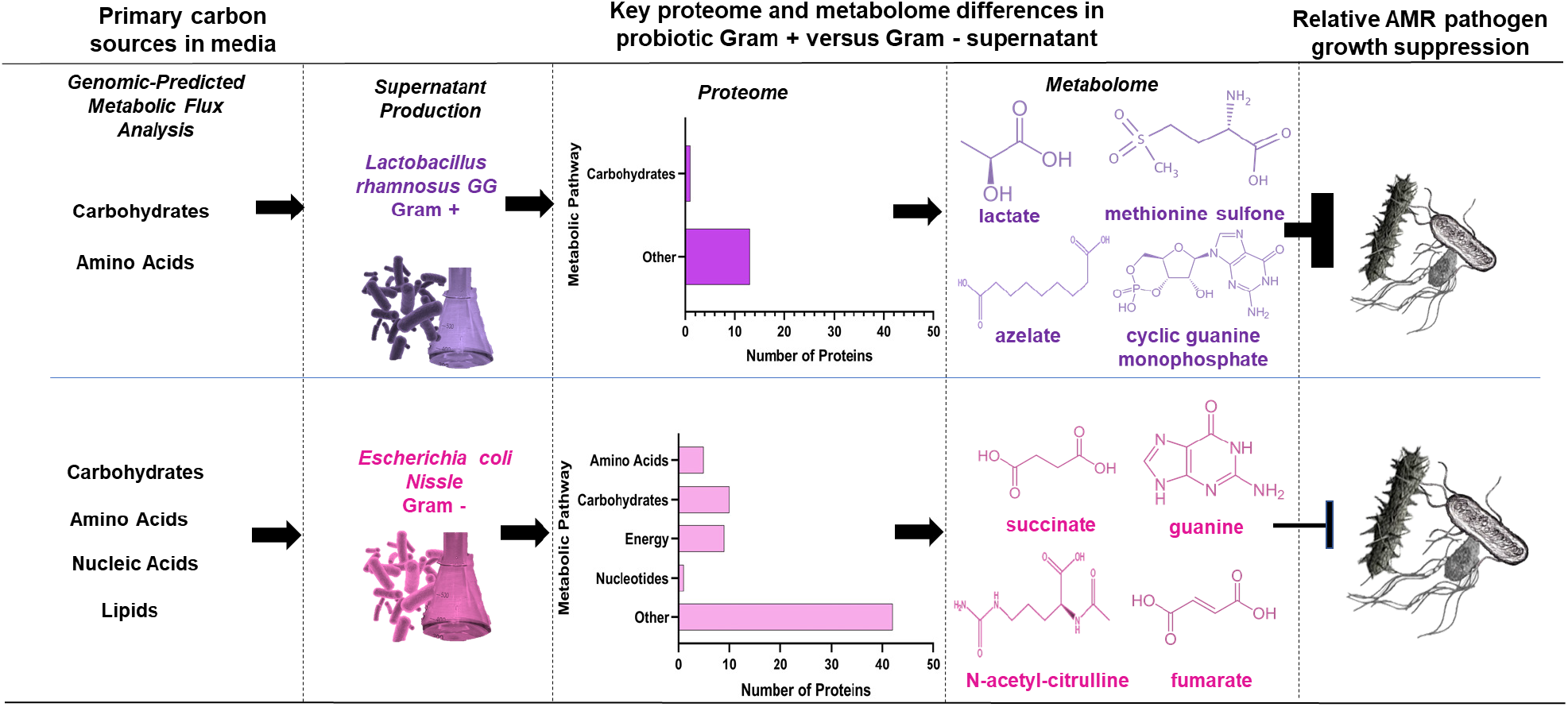
Protein and metabolite profile summary of LGG and ECN probiotic cell free supernatants with distinct efficacy for growth suppression of three AMR pathogens. Metabolic models predicted that utilization of distinct carbon sources was the mechanism for observed differences between the metabolome and proteome of ECN and LGG cell free supernatants. Abbreviations: Antimicrobial Resistance (AMR), cyclic guanosine monophosphate (cGMP), *Escherichia coli* Nissle (ECN), G+ (Gram-positive), G− (Gram-negative), *Lactobacillus rhamnosus* GG (LGG), trimethylamine N-oxide (TMAO).

Carbohydrate metabolism represents a collection of pathways necessary for the generation of ATP through central metabolism to form biosynthetic precursors required for various cellular processes [33]. Utilization of these carbon sources (including amino acid and fatty acid catabolism) differs between Gram-negative and Gram-positive bacteria accounting for the differences in the metabolome and proteome of these probiotics (**Fig. 6**). The metabolic predictive model showed the diversity of metabolic pathways by which ECN was predicted to utilize glycolytic metabolites when compared to LGG, which exhibited more restricted funneling into glycolytic processes. ECN was predicted to expend more energy than LGG to funnel glycolytic metabolites into the synthesis of fatty acids, sugars, and nucleotides, whereas LGG primarily relies on glycolytic metabolites to produce ATP during fermentation reactions (**Fig. 5**). Although diverse glycolytic metabolite shunting observed in ECN provided substrates for other key areas of metabolism, the decreased production of fructose 1,6 diphosphate, glucose 6-phosphate, phosphoenolpyruvate and lactate compared to LGG, may have contributed to the lower bacteriostatic and bactericidal activities of ECN supernatant against Gram-negative bacteria [32].

The presence of type I glyceraldehyde 3-phosphate dehydrogenase in the LGG proteome but not the ECN proteome may additionally contribute to the differential antimicrobial activity of LGG versus ECN supernatant. Interestingly, it was shown that some of the proteins in the glycolytic pathway were localized on the cell wall in some Gram-positive bacteria [34] with the capacity to produce ATP on the cell’s surface [35]. Glyceraldehyde 3-phosphate dehydrogenase (**GAPDH**), which was present in the LGG but not ECN proteome, is one of these proteins (**Fig. 4B**). Previous evaluations of the LGG proteome have shown that GAPDH is not strictly cytosolic, but like with other Gram-positive species, it is secreted into the extracellular environment [36]. In addition to increasing the glycolytic capacity of LGG, GAPDH has been increasingly explored in prokaryotic as well as eukaryotic species to produce antimicrobial peptides that suppress the growth of various Gram-negative pathogens [36]. While probiotic secreted GAPDH has been implicated in host adhesion as well as immunomodulation [37], it has not yet been screened for production of antimicrobial peptide products in probiotic bacteria, and warrants further attention as a LGG mediator of pathogen growth suppression. This hypothesis is consistent with results from integrated metabolic modeling, whereby proteomic and metabolomic observations support differences in LGG and ECN for carbohydrate metabolism. Notably, LGG carbohydrate metabolism simultaneously increased bacteriostatic organic acid production, and LGG secreted GAPDH produced uncharacterized antimicrobial proteins contributing to pathogen growth suppression. (**Fig. 4C)**.

Twenty-six of the 37 metabolites predicted to be consumed by ECN were involved in purine metabolism (**File S3**). *Escherichia coli* has been shown to utilize purines, including guanine, as nitrogen sources and convert exogenous purines (bases or nucleosides) to nucleotides, which are converted to nucleobases [38]. The purine nucleobases are then converted to the corresponding purine mononucleotides by the exo-enzymes hypoxanthine and guanine phosphoribosyltransferase, which salvage guanine, hypoxanthine, and xanthine, three metabolites shown to have antimicrobial activity [39]. Given the lack of nucleotide metabolism proteins in the ECN supernatant, improved recovery and prediction of bioactive proteins secreted into probiotic supernatants is an area for future investigation.

While the collective supernatant metabolomes and proteomes did share metabolites, the supernatant metabolome were unique to function in ECN or LGG. For ECN, exclusive metabolites included N-acetylcitrulline, a metabolite of urea cycle, arginine and proline metabolism [40], dihomo-linoleate (20:2n6) a product of polyunsaturated fatty acid metabolism shown to have antioxidant properties [41], nicotinamide ribose [42], and 3-hydroshikimate a product of the shikimate pathway whose enzymes are targets for the design of potential antimicrobial agents [43]. In *E. coli*, arginine metabolism distinguished probiotic *E. coli* strains from commensal and pathogenic *E. coli* strains, whereas *E. coli* Nissle was shown to produce higher levels of citrulline, citrulline derivatives, and an overall greater diversity of arginine-derived metabolites [20]. The unique production of N-acetylcitrulline and other arginine metabolites, including the proteins and enzymes regulating ECN arginine metabolism thus represents another research mechanistic dimension to optimize the antimicrobial activity of ECN. LGG supernatant exclusive metabolites included cysteine s-sulfate that is produced by the reaction of inorganic sulfite and cystine and a very potent N-methyl-D-aspartate-receptor (NMDA-R) agonist [44], N1-methyladenosine, which plays a role in environmental stress, ribosome biogenesis and antibiotic resistance [45], and nicotinamide adenine dinucleotide (NAD+), a cofactor that is central to metabolism involved in redox reactions. Three unknown metabolites were uniquely detected in LGG. Collectively, the roles for unknown/unnamed metabolites in LGG warrant additional evaluation for metabolic functional relevance to the probiotic, and how production can be increased for antimicrobial applications. Further investigation and quantitation of these metabolites is warranted to characterize these compounds, as they could be contributing to the enhanced growth suppressing effect of LGG supernatant observed when compared to ECN supernatant.

This proteomic, metabolomic and metabolic flux analysis of two diverse probiotic cell free supernatants highlighted the role for carbohydrate, amino acid, and nucleotide metabolism as strategies for suppressing AMR pathogen growth. Our findings herein contributed novel mechanistic insights to the metabolic pathway synergy inferred from in vivo studies that utilize and test ECN and LGG in combination and with prebiotics that enhanced probiotic functions [18, 46, 47]. Targeted quantification and stochiometric evaluation of antimicrobials in cell free supernatants are needed for confirming minimum inhibitory concentrations that suppress pathogen growth, and for impacts on host gastrointestinal and mucosal immune functions.

Optimization of Gram-positive and Gram-negative probiotics for antimicrobial therapies to AMR pathogens requires attention to several environmental and host conditions that allow pathogens to spread and prior to concerns for outbreak infections that may affect animal and human health.

## Materials and Methods

### Antimicrobial resistant pathogen isolation

The *Salmonella enterica* serovar Typhimurium isolate used in this study was collected from human intestinal tract in 2010, Washington State University, and was provided as a generous gift from Dr. Sangeeta Rao at Colorado State University. The AMR *E. coli* and *K. oxytoca* isolates were collected from environmental water samples in Northern Colorado using published methods [10]. Briefly, water samples were collected with sterile Pyrex wide-mouth storage bottles, immediately placed on ice, and kept in a light-sensitive container until analysis, which occurred approximately 1h following sample collection. Water samples were diluted onto CHROMagar-ESBL (extended-spectrum beta-lactamase) and CHROMagar-KPC (*Klebsiella pneumoniae* carbapenemase) (DRG Diagnostics, Springfield, NJ) media to identify and isolate individual colonies. Isolated colonies were incubated in tryptic soy broth (**TSB**) at 37°C for ~18h, and colony identities were made to the species-level using matrix-assisted laser desorption-ionization time-of-flight analysis (**MALDI**) on a VITEK-MS machine (Biomerieux, Durham, NC).

### *Antimicrobial resistance profile determination for Salmonella* Typhimurium, *E. coli*, and *K. oxytoca*

The AMR profiles of *Salmonella* Typhimurium, *E. coli*, and *K. oxytoca* were established using Kirby-Bauer Disc Diffusion methods established by the Clinical & Laboratory Standards Institute (**CLSI**) [48]. Briefly, overnight incubations of each isolate cultured in sterile TSB were diluted to a concentration of 1.5×10^8^ cells/mL using a 0.5 McFarland Standard. The resultant dilutants were spread onto Mueller-Hinton agar (Hardy Diagnostics, Santa Maria, CA) and the following antimicrobial discs were applied: Meropenem (MEM-10), Linezolid (LZD-30), Vancomycin (VA-30), Cefazolin (CZ-30), Ciprofloxacin (CIP-5), Gentamicin (CN-10). Ampicillin (AMP-10), Penicillin (P-10), Tobramycin (NN-10), Tetracycline (TE-30), and Amikacin (AK-30). After 18h incubation at 37°C, the zone of inhibition was measured and reported as the radius from the center of the disc to the edge of the inhibition zone (mm). Kirby-Bauer Disc Diffusion assays were performed in triplicate for each pathogen, and the zone of inhibition was averaged across assays. These averaged antimicrobial disc inhibition zones were compared to CLSI standards for each isolate to make the determinations of “Susceptible”, “Intermediate”, and “Resistant”.

### Whole genome sequencing

DNA was extracted from each isolate using a DNeasy PowerSoil Kit (Qiagen, Valencia, CA) following manufacturer protocols. Extracted DNA was semi-quantified and quality-checked using a NanoDrop 2000 (Thermo Scientific, Lafayette, CO). To confirm sample sterility, sterile TSB broth and DNA extraction media from the DNeasy kit were used as negative controls during extraction and quantitation. Following extraction and quantitation, all samples were stored at −20°C until further analysis.

Extracted DNA samples were sequenced at the South Dakota State University Animal Disease Research and Diagnostic Laboratory by Dr. Joy Scaria and Dr. Linto Antony using previously described methods [49]. Briefly, samples were processed using a Nextera XT DNA Sample Prep Kit (Illumina Inc., San Diego, CA), were subsequently pooled in equimolar amounts, and sequenced on an Illumina Miseq platform (Illumina Inc., San Diego, CA). A 2×250 paired-end approach with V2 chemistry was used to sequence samples. The genome of each sample was assembled using Geneious Prime Version 2019.2.1 (Biomatters Ltd., Auckland, New Zealand) using reference genomes for *E. coli*, *S.* Typhimurium, and *K. oxytoca* made publicly available via the National Center for Biotechnology Information. Each assembled genome was processed through the basic local alignment search tool (**BLAST**) through the MEGARes Database [50] for AMR genes. Positive gene identifies were defined as having 85% sequence similarity over 50% of the sequence when compared to the database sequence.

### Probiotic cultures and cell-free supernatant preparation

The *E. coli* Nissle 1917 and *L. rhamnosus* GG ATCC 53103 isolates used for the experiments herein were provided by Dr. Lijuan Yuan at the Virginia Polytechnic Institute and State University. Cell-free supernatant was prepared as described previously [51].

Approximately 1X10^7^ colony forming units (**CFU**) of each probiotic isolate was propagated in deMan Rogasa Sharpe (**MRS**) broth (Beckton, Dickinson and Company, Difco Laboratories, Franklin Lakes, NJ) for 24h at 37°C. The resultant cultures were centrifuged at 4000xg for 10 minutes, and the supernatant was decanted from the resultant cellular pellet. The supernatant was then centrifuged and decanted again using the same conditions as the initial round and titrated to a pH of 4.50 using a 1 mol*L^−1^ solution of NaOH (Sigma Aldrich, St. Louis, MO) with a pH meter (Corning Pinnacle 530, Cole-Parmer, Vernon Hills, IL). All titrated supernatant was filtered through a 0.22 μM-pore filter (Pall Corporation LifeSciences, Port Washington, NY) before being stored at −80°C prior to use. Three independently prepared (biological replicates) of supernatant were prepared and used in the subsequent analyses described herein.

### Pathogen growth assays and probiotic cell-free supernatant treatments

*S.* Typhimurium, *E. coli*, and *K. oxytoca* isolates were thawed and grown in the presence of probiotic cell-free supernatant as described previously [51]. Frozen −80°C stocks of each pathogen were thawed and grown to early/mid exponential phase using a Cytation3 plate reader (BioTek Instruments Inc., Winooski, VT) and approximately 2×10^5^ CFU/mL of pathogen was inoculated into 180 μL of sterile Luria Bertani (**LB**) broth in a 96-well plate. The following concentrations of cell-free supernatant from LGG and ECN were added to wells inoculated with pathogen: 25% v/v (60 μL), 22% v/v (50 μL), 18% v/v (40 μL) and 12% v/v (25 μL). These supernatant concentrations were guided by previous dose-dependent treatments to *S. enterica* serovar Typhimurium strain 14028s [51]. Equivalent concentrations of sterile MRS, pH 4.50, and sterile LB, pH 4.50, were used as a vehicle control and negative control respectively for each cell-free supernatant treatment. Pathogen growth in the presence of cell-free supernatant was measured every 20 minutes for 18h on a Cytation3 plate-reader using optical density read at a wavelength of 600 nm (**OD600**). To quantify growth suppression at each timepoint, percent growth suppression was calculated by comparing pathogen growth in the presence of a supernatant treatment versus the vehicle control using the following equation:

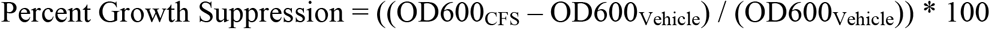

For each pathogen, the growth suppression assay was repeated a minimum of 3 times, and each assay contained a minimum of three technical replicates of each probiotic supernatant concentration. A repeated measures two-way analysis of variance was used to compare treatment optical densities at each time point and p-values were adjusted using a Tukey post-test to control for multiple comparisons. A p-value of p<0.05 was defined as statistically significant. Each supernatant concentration was compared between LGG and ECN for each pathogen (e.g. 25% LGG vs ECN CFS for *S.* Typhimurium, *E. coli*, or *K. oxytoca*).

### Probiotic Cell Free Supernatant Metabolomics

To establish the small molecule profiles of *L. rhamnosus* GG and *E. coli* Nissle cell-free supernatants, the global, non-targeted metabolome of each was determined by Metabolon Inc © (Durham, NC) using previously described methods [51]. Three replicates each of LGG and ECN supernatant, representing three independent supernatant collections, and three replicates of sterile MRS broth were sent to Metabolon on dry ice and stored in liquid nitrogen. Prior to extraction, the protein content of each sample was removed using an 80% ice-cold (−80°C) methanol aqueous solution coupled with vigorous shaking for two minutes and subsequent centrifugation at 680xg for 3 minutes. The resultant samples were each divided into five parts for analysis using ultra-high-performance liquid-chromatography tandem mass-spectrometry (**UPLC-MS/MS**) and consisted of: two aliquots for reverse phase UPLC-MS/MS analysis with positive ion mode electrospray ionization (**ESI**), one aliquot for reverse phase UPLC-MS/MS analysis with negative ion mode ESI, one aliquot for hydrophilic interaction (**HILIC**)/UPLC-MS/MS with negative ion mode ESI, and one backup aliquot. Each aliquot was evaporated using a TurboVap^®^ solvent evaporation system (Zymark, Hopkinton, MA) to remove organic solvent and stored under nitrogen before subsequent analysis.

For UPLC processing, each sample was injected into a Waters ACQUITY UPLC column using solvents optimized for the five aliquot run analyses described above. For the reverse phase UPLC-MS/MS with positive ion mode ESI analysis, one aliquot of each sample was gradient-eluted using a C18 column (Waters UPLC BEH C18-2.1×100mm, 1.7μm) with a water and methanol mobile phase containing 0.05% v/v perfluoropentanoic acid and 0.1% v/v formic acid. A second aliquot for analysis using UPLC-MS/MS with positive ion mode ESI was gradient-eluted using the afore-mentioned C18 column with a mobile phase of methanol, acetonitrile, water, 0.05% v/v perfluoropentanoic acid, and 0.01% formic acid. For the reverse phase UPLC-MS/MS analysis with negative ion mode ESI, an aliquot of each sample was gradient-eluted using a separate C18 column with a mobile phase of methanol, water, and 6.5mM of ammonium bicarbonate at a pH of 8.0. HILIC-UPLC-MS/MS with negative ion mode ESI for each sample was performed on a HILIC column (Waters UPLC BEH Amide 2.1×150 mm, 1.7μm) with a mobile gradient of water, acetonitrile, and 10mM of ammonium format at a pH of 10.8.

Following gradient elution, all samples were subjected to MS/MS processing using a Thermo Scientific Q-Exactive high resolution/accurate mass spectrometer operating with an Orbitrap mass analyzer at 35,000 mass resolution and coupled with heated electrospray ionization source. Mass spectral scans utilized dynamic exclusion with both MS and data-dependent MS^n^ scans to detect peaks, and the scan range covered approximately 70-1000 m/z. Raw data from MS/MS scans were peak-identified and processed for quality control using proprietary Metabolon .NET systems that compared data to known sources of artifacts and background noise inherent to each UPLC-MS/MS run type. Peak identification was made by comparing peaks to an internal database of ~3,300 purified chemical standards using retention indices, m/z ratios (within +/− 10 ppm of the purified internal standards), and MS/MS forward and reverse match scores. Peaks that did not match a purified internal standard but had a retention index and m/z that was not artifact or background noise were reported as “unknown” in subsequent analyses.

### Metabolite normalization, statistical analysis, and visualization

The raw abundances for detected metabolite were normalized using area under the curve analysis, where the raw abundance of each metabolite was divided by the median raw abundance of the metabolite across the dataset. One-way analysis of variance was used to compare the median-scaled abundance of each metabolite across sample types, where statistical significance was defined as a p <0.05. To account for false positive detections, q-values were calculated for each metabolite, and metabolites with a q>0.1 were excluded from downstream analysis. Pathway enrichment scores (**PES**) were calculated to evaluate the contribution of different metabolic pathways to overall metabolite profile differences between treatments using the following formula, where “k” indicates the number of significant metabolites in the metabolic pathway, “m” indicates the total number of metabolites in the metabolic pathway, “n” indicates the number of significant metabolites in the entire data set, and “N” indicates the total number of metabolites in the data set.

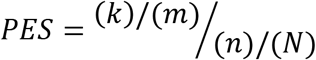

Median-scaled abundances were additionally used to calculate metabolite fold differences by dividing the average median-scaled abundance of a metabolite in one sample by its average median scaled abundance in a second sample type (e.g. average metabolite abundance in ECN supernatant versus average metabolite abundance in LGG supernatant). Metabolite visualization was performed using Metaboanalyst^®^ (version 4.0) with R version 3.6.1, using the raw abundance of each metabolite generated by Metabolon [52, 53] (R-script **File S6**). A principal coordinates analysis plot was generated using metabolite median-scaled abundances. A heat map with hierarchical clustering analysis was generated using Euclidean and Ward differential clustering algorithms, where red boxes indicate metabolites that were elevated in ECN compared to LGG, and blue boxes indicate metabolites that were decreased in ECN versus LGG The heat map visualizes the fifty metabolites with the largest statistical differences when comparing ECN versus LGG.

### Probiotic cell free supernatant proteomics

Non-targeted proteomes of each probiotic supernatant and sterile growth media (MRS) were generated by the Colorado State University Proteomics and Metabolomics Facility using LC-MS/MS. Proteins were isolated from supernatant in a 1:4 v/v suspension of ice-cold (−80°C) methanol. The resultant protein pellets were washed in 100% ice-cold (−80°C) acetone and centrifuged at 15,000xg for 10 minutes. Following two rounds of acetone washing, samples were air-dried and reconstituted in 2M urea and bath sonicated for five minutes. To collect insoluble material, sonicated samples were centrifuged at 4000xg for two minutes. To quantitate samples, aliquots were diluted 1:2 and 1:5 in 2M urea solvent and their total protein concentration was measured using a Pierce Bicinchoninic Acid Protein Assay (Thermo Scientific, Waltham, MA) following manufacturer’s instructions. Approximately 50 μg of each sample was subjected to trypsin digestion using the methods described by Schauer *et al.* 2013 [54]. Briefly, 50 μg of each sample was reconstituted in a solution containing 8M urea, 0.2% v/v ProteaseMax ™ surfactant trypsin enhancer (Promega, Madison, WI), 5mM dithiothreitol, and 5mM iodoacetic acid.

Purified trypsin (Pierce MS-Grade, Thermo Scientific, Waltham, MA) was added at a 1:28 ratio to the sample proteins, and the slurry was incubated at 37°C for 3h, after which trypsin was deactivated using 5% trifluoroacetic acid. Desalting occurred using Pierce C18 spin columns following manufacturer instructions (Thermo Scientific, Waltham, MA). The eluates were dried in a vacuum evaporator and reconstituted in 5% v/v acetonitrile and 0.1% v/v formic acid. Total peptide quantification was determined for each sample resuspension on a NanoDrop (Thermo Scientific, Waltham, MA) at a wavelength of 205nm and normalized using an extinction coefficient of 31 [55].

Reverse phase chromatography was performed using water with 0.1% formic acid and acetonitrile with 0.1% formic acid. A total of 0.75μg of peptides was purified and concentrated using an on-line enrichment column (Waters Symmetry Trap C18 100Å, 5μm, 180 μm ID x 20mm column). Subsequent separation was performed using a reverse-phase C18 nanospray column (Waters, Peptide BEH C18; 1.7μm, 75μm x 150nm column) at 45°C and samples were eluted using a 30-minute mobile phase gradient of 3-8% formic acid over 3 minutes, followed by 8%-35% of a acetonitrile with 0.1% formic acid solution over 27 minutes, at a flow rate of 350 nL/min. A Nanospray Flex ion source (Thermo Scientific, Waltham, MA) introduced eluate directly into the mass spectrometer (Orbitrap Velos Pro™, Thermo Scientific, Waltham, MA).

Spectra were collected using positive ion mode over a range of 400-2,000 m/z, and MS/MS was performed on ions assigned a charge state of 2+ or 3+ using a dynamic exclusion limit of 2 MS/MS spectra of a given m/z value for 30 s (exclusion duration of 90 s). Fourier-Transformation mode (60,000 resolution) was applied for MS detection, and ion trap mode was applied for the subsequent MS/MS with 35% normalized collision energy. Compound lists of the resulting spectra were generated using Xcalibur 3.0 software (Thermo Scientific) with a S/N threshold of 1.5 and 1 scan/group.

### Proteome identification and normalization

MS/MS spectra for each sample were extracted, charge state deconvoluted and deisotoped by ProteoWizard MsConvert (version 3.0). All spectra were then screened for protein identities using Mascot (Matrix Science, London, UK, version 2.6.0) with a fragment ion mass tolerance of 0.80 Daltons and a parent ion tolerance of 20 ppm. Carboxymethylation of cysteine was specified in Mascot as a fixed modification. Deamidation of asparagine and glutamine, methylation of lysine and arginine, hydroxylation of proline, oxidation of methionine, dimethylation of lysine and arginine and acetylation of the n-terminus were specified in Mascot as variable modifications. The following reverse concatenated Uniprot reference proteomes were used for the search: Uniprot_Yeast_rev_022119, Uniprot_Sus_scrofa_rev_022119, Uniprot_Bovine_rev_022119. LGG supernatant samples were additionally screened with the Uniprot_Lactobacillus_rhamnosus_GG_rev_021819 database and ECN supernatant samples were also screened with the Uniprot_Escherichia_coli_Nissle_rev_021819 database. Identified spectra were further combined using the probabilistic protein identification algorithms [56] utilized by Scaffold (version 4.8.4, Proteome Software Inc., Portland, OR) [57]. The peptide probability threshold was set (90%) such that a peptide false discovery rate of 0.0% was achieved based on hits to the reverse database [58]. Protein identifications were accepted if they could be established at greater than 95.0% probability as assigned by the Protein Prophet algorithm [67] and contained at least two identified peptides. Proteins that contained similar peptides and could not be differentiated based on MS/MS analysis alone were grouped to satisfy the principles of parsimony. For each identified protein, raw abundances were used to derive the normalized abundance factor (NSAF) within each sample, a method used to estimate the protein content within a single sample or gel band. NSAF is calculated using the number of spectra (SpC) identifying a protein divided by the protein length (L), referred to as Spectral Abundant Factor (SAF) and then normalized over the total sum of spectral counts/length in a given analysis.

### Metabolic modeling of E. coli Nissle and L. rhamnosus GG

Draft metabolic models for ECN and LGG were reconstructed using apps in DoE KBase [59] iML1515, a published metabolic model for *E. coli* K-12 MG1655 [60], was used as a reference model for ECN. A model for *L. casei* ATCC 344 [61] was used as a reference model for LGG. The genomes for the four organisms were accessed through the KBase interface for NCBI genomes. The KBase application ‘Compare Two Proteomes’ was used to find orthologs between the strain, and the application ‘Propagate Model to New Genome’ was then used to translate the model for *E. coli* K-12 to ECN and the model for *L. casei* ATCC 344 to LGG, respectively, provided with the ortholog comparisons, and it also performed gap-filling to ensure biomass production. Further gap-filling was performed manually to include pathways for the consumption and production of the metabolites detected in the supernatant metabolome. To simulate flux distributions consistent with the supernatant metabolome data for ECN and LGG, an optimization problem was constructed to solve parsimonious flux balance analysis (**pFBA**) [62] for both models simultaneously. Within the models, variables controlled for included metabolite availability in the MRS media and uptake/export of a metabolite by ECN or LGG based on the relative abundance from the metabolome data. Ten thousand flux distributions were simulated under randomly sampled maximum substrate uptake (which were not measured during the experiments) using pFBA maximization of biomass production in the models as the objective function [63]. **File S7** provides the complete mathematical formulation used to construct this modeling analysis. The shadow price for each metabolite was retrieved from the solution of the optimization problem. All simulations were performed in MATLAB R2017b using the COBRA toolbox [64] and the optimization solvers GUROBI [65] and IBM CPLEX [66].

### Data availability

The whole genome sequences used in this study are fully accessible for download at the National Center of Biotechnology Information (NCBI) Sequence Read Archive (SRA) via the following link (Accession Number: PRJNA530250): https://www.ncbi.nlm.nih.gov/bioproject/PRJNA530250

The complete raw data and R script for metabolomics analysis, Metaboanalyst visualization, as well as raw proteome data are provided as supplemental data files. The files for the metabolic modeling analysis are available at https://github.com/chan-csu/modelEcnLggExoMetabolomes’

## Declarations

The authors declare no conflicts of interest.

## Acknowledgements

The authors would like to thank Dr. Sangeeta Rao, PhD (Colorado State University) for donating the *S.* Typhimurium isolate and Dr. Lijuan Yuan, PhD (Virginia Polytechnic Institute) for donating the *E. coli* Nissle and *L. rhamnosus* GG isolates used in this study. We also acknowledge the technical assistance from Dr. Corey Broeckling, Dr. Lisa Wolfe and Kitty Brown for proteomics methods development and data processing at the Colorado State University Analytical Resources Core: Bioanalysis and Omics (ARC-BIO). Additional thanks to Dr. Joy Scaria and Dr. Linto Antony for sequencing pathogen genomes at the South Dakota State University Animal Disease Research & Diagnostic Laboratory. The authors would like to acknowledge funding support from the Bill and Melinda Gates Foundation (OPP1043255) and the National Institutes of Health-Ruth L. Kirschstein-National Research Service Program (5T32OD012201-05).

## Abbreviations

AMR: Antimicrobial Resistance
BLAST: Basic Local Alignment Search Tool
CFS: Cell-Free Supernatant
CFU: Colony Forming Unit
CLSI: Clinical and Laboratory Standards Institute
ECN: *Escherichia coli* Nissle
ESI: Electrospray Ionization
GAPDH: Glyceraldehyde-3-Phosphate Dehydrogenase
HILIC: Hydrophilic Interaction Liquid Chromatography
LB: Luria Bertani
LC-MS/MS: Liquid Chromatography-Tandem Mass Spectrometry
LGG: *Lactobacillus rhamnosus GG*
m/z: Mass to Charge Ratio
MALDI: Matrix-Assisted Laser Desorption/Ionization
MRS: deMan Rogasa Sharpe
NAD+: Nicotinamide Adenine Dinucleotide (oxidized)
OD600: Optical Density, 600-nanometer wavelength
PES: Pathway Enrichment Score
pFBA: Parsimonious Flux-Based Analysis
RI: Retention Index
TCA: Tricarboxylic Acid Cycle
TSB: Tryptic Soy Broth
UPLC-MS/MS: Ultra-High-Performance Liquid Chromatography-Tandem Mass Spectrometry

## References

1. Sanders, M.E., et al., Shared mechanisms among probiotic taxa: implications for general probiotic claims. Curr Opin Biotechnol, 2018. 49: p. 207–216.

2. Sniffen, J.C., et al., Choosing an appropriate probiotic product for your patient: An evidence-based practical guide. PloS one, 2018. 13(12): p. e0209205–e0209205.

3. Liévin-Le Moal, V. and A.L. Servin, Anti-infective activities of lactobacillus strains in the human intestinal microbiota: from probiotics to gastrointestinal anti-infectious biotherapeutic agents. Clinical microbiology reviews, 2014. 27(2): p. 167–199.

4. Ansari, J.M., et al., Strain-level diversity of commercial probiotic isolates of Bacillus, Lactobacillus, and Saccharomyces species illustrated by molecular identification and phenotypic profiling. PloS one, 2019. 14(3): p. e0213841–e0213841.

5. Liu, J., et al., Lactobacillus plantarum ZS2058 and Lactobacillus rhamnosus GG Use Different Mechanisms to Prevent Salmonella Infection in vivo. Frontiers in microbiology, 2019. 10: p. 299–299.

6. Kandasamy, S., et al., Differential Effects of Escherichia coli Nissle and Lactobacillus rhamnosus Strain GG on Human Rotavirus Binding, Infection, and B Cell Immunity. Journal of immunology (Baltimore, Md. : 1950), 2016. 196(4): p. 1780–1789.

7. Magrini, E.T.a.N., Global Priority List of Antibiotic-Resistant Bacteria to Guide Research, Discovery, And Development of New Antibiotics. 2018, World Health Organization.

8. Antibiotic Resistance Threats in the United States, U.S.D.o.H.a.H. Services, Editor. 2019, Centers for Disease Control and Prevention: Athens, GA. USA.

9. Salmonella (non-typhoidal). 2018, World Health Organization.

10. Haberecht, H.B., et al., Antimicrobial-Resistant Escherichia coli from Environmental Waters in Northern Colorado. Journal of environmental and public health, 2019. 2019: p. 3862949–3862949.

11. Wangchinda, W., et al., Collateral damage of using colistin in hospitalized patients on emergence of colistin-resistant Escherichia coli and Klebsiella pneumoniae colonization and infection. Antimicrobial resistance and infection control, 2018. 7: p. 84–84.

12. Pitout, J.D.D., P. Nordmann, and L. Poirel, Carbapenemase-Producing Klebsiella pneumoniae, a Key Pathogen Set for Global Nosocomial Dominance. Antimicrobial agents and chemotherapy, 2015. 59(10): p. 5873–5884.

13. Lake, J.G., et al., Pathogen Distribution and Antimicrobial Resistance Among Pediatric Healthcare-Associated Infections Reported to the National Healthcare Safety Network, 2011–2014. Infect Control Hosp Epidemiol, 2018. 39(1): p. 1–11.

14. Vuotto, C., et al., Antibiotic Resistance Related to Biofilm Formation in Klebsiella pneumoniae. Pathogens (Basel, Switzerland), 2014. 3(3): p. 743–758.

15. Naik, A.K., et al., Lactobacillus rhamnosus GG reverses mortality of neonatal mice against Salmonella challenge. Toxicology research, 2019. 8(3): p. 361–372.

16. He, X., et al., Lactobacillus rhamnosus GG supernatant enhance neonatal resistance to systemic Escherichia coli K1 infection by accelerating development of intestinal defense. Scientific reports, 2017. 7: p. 43305–43305.

17. Kandasamy, S., et al., Unraveling the Differences between Gram-Positive and Gram-Negative Probiotics in Modulating Protective Immunity to Enteric Infections. Frontiers in immunology, 2017. 8: p. 334–334.

18. Nealon, N.J., et al., Rice Bran and Probiotics Alter the Porcine Large Intestine and Serum Metabolomes for Protection against Human Rotavirus Diarrhea. Frontiers in Microbiology, 2017. 8(653).

19. Shah, P., et al., A microfluidics-based in vitro model of the gastrointestinal human-microbe interface. Nature communications, 2016. 7: p. 11535–11535.

20. van der Hooft, J.J.J., et al., Substantial Extracellular Metabolic Differences Found Between Phylogenetically Closely Related Probiotic and Pathogenic Strains of Escherichia coli. Frontiers in microbiology, 2019. 10: p. 252–252.

21. Engevik, M.A. and J. Versalovic, Biochemical Features of Beneficial Microbes: Foundations for Therapeutic Microbiology. Microbiology spectrum, 2017. 5(5): p. 10.1128/microbiolspec.BAD-0012-2016.

22. Revelles, O., et al., The carbon storage regulator (Csr) system exerts a nutrient-specific control over central metabolism in Escherichia coli strain Nissle 1917. PloS one, 2013. 8(6): p. e66386–e66386.

23. Manuel, J., G.G. Zhanel, and T. de Kievit, Cadaverine Suppresses Persistence to Carboxypenicillins in *Pseudomonas aeruginosa* PAO1. Antimicrobial Agents and Chemotherapy, 2010. 54(12): p. 5173–5179.

24. Cagno, V., et al., The Agmatine-Containing Poly(Amidoamine) Polymer AGMA1 Binds Cell Surface Heparan Sulfates and Prevents Attachment of Mucosal Human Papillomaviruses. Antimicrobial Agents and Chemotherapy, 2015. 59(9): p. 5250–5259.

25. Su, R.B., et al., Antimalarial effect of agmatine on Plasmodium berghei K173 strain. Acta Pharmacol Sin, 2003. 24(9): p. 918–22.

26. Shukla, S., et al., Detection of biogenic amines and microbial safety assessment of novel Meju fermented with addition of Nelumbo nucifera, Ginkgo biloba, and Allium sativum. Food Chem Toxicol, 2018. 119: p. 231–236.

27. Schroll, C., et al., Polyamines are essential for virulence in Salmonella enterica serovar Gallinarum despite evolutionary decay of polyamine biosynthesis genes. Vet Microbiol, 2014. 170(1-2): p. 144–50.

28. Shwartzman, G., Concerted antibiotic effect of penicillin, methionine, threonine and methionine sulfoxide upon brucella, eberthella, salmonella, and shigella. Science, 1945. 102(2641): p. 148–50.

29. Brenchley, J.E., Effect of methionine sulfoximine and methionine sulfone on glutamate synthesis in Klebsiella aerogenes. J Bacteriol, 1973. 114(2): p. 666–73.

30. Hentchel, K.L. and J.C. Escalante-Semerena, In Salmonella enterica, the Gcn5-related acetyltransferase MddA (formerly YncA) acetylates methionine sulfoximine and methionine sulfone, blocking their toxic effects. J Bacteriol, 2015. 197(2): p. 314–25.

31. Krajewski, S.S., I. Isoz, and J. Johansson, Antibacterial and antivirulence effect of 6-N-hydroxylaminopurine in Listeria monocytogenes. Nucleic acids research, 2017. 45(4): p. 1914–1924.

32. Stanojević-Nikolić, S., et al., Antimicrobial Activity of Lactic Acid Against Pathogen and Spoilage Microorganisms. Journal of Food Processing and Preservation, 2016. 40(5): p. 990–998.

33. Richardson, A.R., G.A. Somerville, and A.L. Sonenshein, Regulating the Intersection of Metabolism and Pathogenesis in Gram-positive Bacteria. Microbiology spectrum, 2015. 3(3): p. 10.1128/microbiolspec.MBP-0004-2014.

34. Pancholi, V. and V.A. Fischetti, A major surface protein on group A streptococci is a glyceraldehyde-3-phosphate-dehydrogenase with multiple binding activity. Journal of Experimental Medicine, 1992. 176(2): p. 415–426.

35. Fischetti, V.A., Surface Proteins on Gram-Positive Bacteria. Microbiology spectrum, 2019. 7(4): p. 10.1128/microbiolspec.GPP3-0012-2018.

36. Branco, P., et al., Identification of novel GAPDH-derived antimicrobial peptides secreted by Saccharomyces cerevisiae and involved in wine microbial interactions. Appl Microbiol Biotechnol, 2014. 98(2): p. 843–53.

37. Sánchez, B., P. Bressollier, and M.C. Urdaci, Exported proteins in probiotic bacteria: adhesion to intestinal surfaces, host immunomodulation and molecular cross-talking with the host. FEMS Immunology & Medical Microbiology, 2008. 54(1): p. 1–17.

38. Xi, H., B.L. Schneider, and L. Reitzer, Purine catabolism in Escherichia coli and function of xanthine dehydrogenase in purine salvage. Journal of bacteriology, 2000. 182(19): p. 5332–5341.

39. Raj, C.V. and S. Dhala, Effect of naturally occurring xanthines on bacteria. I. Antimicrobial action and potentiating effect on antibiotic spectra. Appl Microbiol, 1965. 13(3): p. 432–6.

40. van der Hooft, J.J.J., et al., Substantial Extracellular Metabolic Differences Found Between Phylogenetically Closely Related Probiotic and Pathogenic Strains of Escherichia coli. Frontiers in Microbiology, 2019. 10(252).

41. Guermouche, B., et al., Effect of dietary n -3 polyunsaturated fatty acids on oxidant/antioxidant status in macrosomic offspring of diabetic rats. BioMed research international, 2014. 2014: p. 368107–368107.

42. Mehmel, M., N. Jovanović, and U. Spitz, Nicotinamide Riboside-The Current State of Research and Therapeutic Uses. Nutrients, 2020. 12(6).

43. Gower, M.A., Synthesis and biological evaluation of inhibitors of the shikimate pathway enzyme 3-dehydroquinate dehydratase. 2006, University of Canterbury.

44. Zecchini, M., R. Lucas, and A. Le Gresley, New Insights into the Cystine-Sulfite Reaction. Molecules, 2019. 24(13).

45. Chen, Y., et al., N1-Methyladenosine detection with CRISPR-Cas13a/C2c2. Chemical Science, 2019. 10(10): p. 2975–2979.

46. Yang, X., et al., High protective efficacy of rice bran against human rotavirus diarrhea via enhancing probiotic growth, gut barrier function, and innate immunity. Sci Rep, 2015. 5: p. 15004.

47. Lei, S., et al., High Protective Efficacy of Probiotics and Rice Bran against Human Norovirus Infection and Diarrhea in Gnotobiotic Pigs. Front Microbiol, 2016. 7: p. 1699.

48. Institute, C.a.L.S., M100 Performance Standards for Antimicrobial Susceptibility Testing, in Zone Diameter and Minimal Inhibitory Concentration Breakpoints for Enterobacteriaceae. 2017, Clinical and Laboratory Standards Institute: Wayne, PA. p. 9.

49. Thomas, M., et al., Whole genome sequencing-based detection of antimicrobial resistance and virulence in non-typhoidal Salmonella enterica isolated from wildlife. Gut pathogens, 2017. 9: p. 66–66.

50. Doster, E., et al., MEGARes 2.0: a database for classification of antimicrobial drug, biocide and metal resistance determinants in metagenomic sequence data. Nucleic acids research, 2020. 48(D1): p. D561–D569.

51. Nealon, N.J., C.R. Worcester, and E.P. Ryan, Lactobacillus paracasei metabolism of rice bran reveals metabolome associated with Salmonella Typhimurium growth reduction. J Appl Microbiol, 2017. 122(6): p. 1639–1656.

52. Chong, J., D.S. Wishart, and J. Xia, Using MetaboAnalyst 4.0 for Comprehensive and Integrative Metabolomics Data Analysis. Current Protocols in Bioinformatics, 2019. 68(1): p. e86.

53. Pang, Z., et al., MetaboAnalystR 3.0: Toward an Optimized Workflow for Global Metabolomics. Metabolites, 2020. 10(5).

54. Schauer, K.L., et al., Proteomic profiling and pathway analysis of the response of rat renal proximal convoluted tubules to metabolic acidosis. Am J Physiol Renal Physiol, 2013. 305(5): p. F628–40.

55. Scopes, R.K., Measurement of protein by spectrophotometry at 205 nm. Anal Biochem, 1974. 59(1): p. 277–82.

56. Keller, A., et al., Empirical statistical model to estimate the accuracy of peptide identifications made by MS/MS and database search. Anal Chem, 2002. 74(20): p. 5383–92.

57. Searle, B.C., M. Turner, and A.I. Nesvizhskii, Improving sensitivity by probabilistically combining results from multiple MS/MS search methodologies. J Proteome Res, 2008. 7(1): p. 245–53.

58. Käll, L., et al., Assigning significance to peptides identified by tandem mass spectrometry using decoy databases. J Proteome Res, 2008. 7(1): p. 29–34.

59. Arkin, A.P., et al., KBase: The United States Department of Energy Systems Biology Knowledgebase. Nature Biotechnology, 2018. 36(7): p. 566–569.

60. Monk, J.M., et al., iML1515, a knowledgebase that computes Escherichia coli traits. Nat Biotechnol, 2017. 35(10): p. 904–908.

61. Vinay-Lara, E., et al., Genome-scale reconstruction of metabolic networks of Lactobacillus casei ATCC 334 and 12A. PloS one, 2014. 9(11): p. e110785–e110785.

62. Orth, J.D., I. Thiele, and B.Ø. Palsson, What is flux balance analysis? Nature Biotechnology, 2010. 28(3): p. 245–248.

63. Lewis, N.E., et al., Omic data from evolved E. coli are consistent with computed optimal growth from genome-scale models. Molecular systems biology, 2010. 6: p. 390–390.

64. Heirendt, L., et al., Creation and analysis of biochemical constraint-based models using the COBRA Toolbox v.3.0. Nat Protoc, 2019. 14(3): p. 639–702.

65. Gurobi Optimizer Reference Manual. 2020, Gurobi Optimization, LLC.

66. Corporation, I.B.M.I., ed. IBM ILOG CPLEX Optimization Studio CPLEX User’s Manual. 12 ed. 2017. 596.

